# Revealing the molecular mechanisms underlying neural transmission of female pheromone signals in the male antennae of *Antheraea pernyi* by integrative proteomics and metabolomics analysis

**DOI:** 10.1101/2022.08.25.505247

**Authors:** Guobao Wang, Xiang Ji, Lei Nie

## Abstract

Detection of sex pheromones of insects relies on the antennae. The female pheromone signal transmission in the male antennae ultimately initiates the courtship and mating behaviors of males. To investigate the proteins and metabolites involved in this neural transduction, the study adopted integrative proteomics and metabolomics analysis including tandem mass tag (TMT) proteomic quantification and liquid chromatography tandem mass spectrometry (LC/MS)-based metabolomics for comparing proteomic and metabolic changes between the antennae of male moths following stimulation by females and the non-stimulated males of *A. pernyi*. A total of 92 differentially expressed proteins (DEPs) containing 52 up-regulated and 40 down-regulated proteins and 545 differentially expressed metabolites (DEMs) including 218 up- and 327 down-regulated metabolites were identified from the antennae of female-stimulated male moths based on the proteome and metabolome data, respectively. GO enrichment analysis showed that 45 DEPs could be enriched into different GO terms on different levels. COG analysis indicated that 61 DEPs were assigned to 20 functional categories. The 160 DEMs respectively fell into 11 and 44 classes at SuperClass and Class levels based on HMDB annotation. KEGG pathway enrichment analysis indicated that totally 43 DEMs were enriched into 6, 27, and 87 pathways on level 1, 2, and 3, respectively. A number of DEPs and DEMs related to neural transmission of female pheromone signals in the male antennae of *A. pernyi* were screened, including tyrosine hydroxylase, cryptochrome-1, tachykinin, arylalkylamine *N*-acetyltransferase, cadherin-23, glutathione *S*-transferase delta 3, tyramine, tryptamine, n-oleoyl dopamine, n-stearoyl dopamine, and n-stearoyl tyrosine. We concluded that the altered expression levels of those proteins or metabolites were involved in regulating the neuron activity for enhanced transmission of neural impulses and continuous perception, reception, and transduction of female pheromone signals. Our findings yielded novel insights into the potential molecular mechanisms in the antennae of male *A. pernyi* responding to female attraction.

## 1. Introduction

Insects perceive multiple environmental chemicals such as volatile and non-volatile odors through chemo-receptors which are situated on olfaction sensilla mainly distributed in the antennae to acquire information about oviposition sites, food resources, social environment, and reproductive partners [1]. The courtship behavior of insects by which an adult male detects and searches for its mate depending on sex pheromones specific to species released by a conspecific female relies on the olfactory function of antennae [2]. In the German cockroach *Blattella germanica*, upon antennal contact with the female body and antennal “fencing”, the male detects the sex pheromones for female contact using their antennae and subsequently execute a wing-raising (WR) display accompanied by a specialized tergal gland exposing to the females for the following copulation [3–5]. A high proportion of male *Teleogryllus commodus* with antennae removal never court conspecific females, not coincidentally, only 45% of the adult male *Gryllus bimaculatus* with both antennae excision courted intact females, suggesting the essential role of antennae in eliciting the male courtship response to females [6]. For *Trichopria drosophilae*, stereotyped and strong antennal courtships that enable two fourth antennomeres in males to contact with two apical antennomeres in females lead to copulation, whereas the mating was restricted by preventing transfer of courtship pheromones from antennae of males to females [7]. Besides, the courtship behavior based on the extensive use of the antennae also has been widely observed in true bugs [8–10], beetles [11], butterflies [12], and hymenopterans [13–15].

During the mates seeking and courtship behavior of male lepidopterous moths, the female pheromones are captured by the male antennae and absorbed via pores on sensilla surfaces. Following bound to the pheromone binding proteins (PBPs), the pheromone-PBP complex passes through the sensillum lymph to activate pheromone receptors expressed on dendritic membranes of the olfactory receptor neurons, which leads to the translation of the chemical signals to electrical signals transducing to the antennal lobe [16–18]. Whereas, there is still a lack of detailed reports on response mechanisms of male moths to female stimulation especially the related proteins and metabolites underlying the female pheromone signal transmission in the male antennae.

Recently, high-throughput sequencing (HTS) technologies have been widely used to explore the olfactory mechanisms of the antennae in various Lepidoptera species such as *Bombyx mori* [19, 20], *Spodoptera frugiperda* [21], *Spodoptera exigua* [22], *Histia rhodope* [23], *Lobesia botrana* [24], *Epiphyas postvittana* [25]. *A. pernyi* is a rising Lepidoptera model organism with several advantages, such as ease rearing and experimental operation [26]. The study applied HTS strategies including LC/MS-based metabolomics and TMT proteomic quantification to explore metabolic and proteomic changes of antennae of male moths displaying wing-flapping (WF males) following stimulation by females compared with the non-stimulated males of *A. pernyi*. The differentially expressed proteins (DEPs) and differentially expressed metabolites (DEMs) were screened. In addition, diverse proteins and metabolites involved in neural transmission of sex pheromone signals in the male antennae undergoing significant alteration upon female stimulation were analyzed. The results are beneficial for better understanding the response mechanisms of the antennae of male *A. pernyi* to female pheromones.

## 2 Materials and methods

### 2.1 Antennae sample collection

The newly emerged adult *A. pernyi* (Xuanda strain) male and female moths were obtained from Liaoning Engineering & Technology Research Center for Insect Resources in Shenyang Agricultural University (Shenyang City, China), kept separately to prevent contact. The male moths were classified into two groups: a control group containing males with mature virgin while no contact with any females and an experimental group containing the males showing wing-flapping (WF males) display following contact with the females for 2-5 min. The complete antennae were dissected from the moths in the two groups and immediately stored at −80 °C.

### 2.2 Protein sample preparation and digestion

Liquid nitrogen and a lysis buffer (LBM-200, Micron biolab) were separately used for powering and homogenizing each antennae sample. After being sonicated on ice, homogenate was subject to 10-min centrifugation under 20000 *g*, followed by filtering and quantification of the supernatant with a BCA kit (BCAK-200, Micron biolab). After reduction using 5 mM dithiothreitol (DTT) for 60 min at 30 °C, proteins of equal amount were alkylated by adopting iodoacetamide (IAM) of 30 mM at room temperature for 1 h in darkness, followed by precipitation using ice-cold acetone. After being washed for three times with acetone and suspension in triethylammonium bicarbonate (TEAB) of 0.1 M, the precipitate experienced digestion using 1/25 trypsin at 37 ℃ for 12 h, which was ended using 1% trifluoroacetic acid (TFA). After cleaning using with Strata X C18 SPE column (Phenomenex), the resultant peptide was then dried in vacuum using a Scanvac maxi-beta (Labogene).

### 2.3 TMT labeling and HPLC fractionation

After reconstituting resultant peptide (2 mg in each sample) in 0.5 M TEAB of 120 μL, it was treated using a TMT10plex label reagent kit (90111, Thermo Scientific) following manufacturing specifications. Then, reversed phase (RP) chromatography was adopted to fractionate labeled peptides through use of a LC20AD HPLC system (Shimadzu) with an XBridge Shield C_18_ RP column (4.6 × 250 mm, 3.5 µm, Waters). The vacuum centrifugation was applied to dry the collected fractions 45 °C.

### 2.4 Proteome analysis by LC-MS/MS

The nanoLC-MS/MS analysis was performed by injecting each fraction. After being loaded in an Acclaim PepMap 100 C_18_ trap column (75 μm × 20 mm, Dionex) by Ultimate 3000 nanoUPLC (Dionex) in solvent A (0.1% FA in H_2_O), peptide mixture was eluted to a home-made RP analytical column (100 μm i.d., 30 cm) containing Reprosil-Pur C_18_ beads of 1.9 μm (Dr. Maisch, Ammerbuch, Germany). This was followed by separation using solvent B of a linear gradient (0.1% FA in ACN) flowing at 450 nL/min. The collected peptide underwent mass spectrometry Orbitrap Exploris 480 (Thermo Scientific) with the nano-spray ionization (NSI) source coupled to UPLC online.

### 2.5 Proteome database search

Maxquant (v.1.5.2.8) was applied for retrieving the resulting raw data. The tandem mass spectrum was searched in the database of *A. pernyi* proteome (https://bigd.big.ac.cn/gwh/, accession number (AN): GWHABGR00000000) [27]. To identify proteins, following options were used: Enzyme used was Trypsin/P; Missed cleavage was 2; false discovery rate (FDR) ≤ 0.01; Precursor mass tolerance was 10 ppm; Fragment mass tolerance was 0.02 Da; Fixed modification included Alkylation (C); Variable modifications included Oxidation (M), Deamidation (NQ), and Acetylation (protein N-term).

For the purpose of comparing protein abundances of different samples, a TMT quantification (TMTQ) algorithm was adopted. By using an FDR (Benjamini-Hochberg), adjustment of p value was realized in many tests. On the basis of analyzing three biological replicates, significantly altered proteins were defined as those, the ratio of the experimental/control TMTQ intensity of which was ≥ 1.2 or ≤ 1/1.2 (p ≤ 0.05). The UniProt-GOA database (https://www.http://www.ebi.ac.uk/GOA/) was adopted to derive Gene Ontology (GO) annotations of proteome. The COG database (Clusters of Orthologous Groups) of proteins (http://www.ncbi.nlm.nih.gov/COG/) was taken to classify functions of the DEPs.

### 2.6 Metabolite extraction and LC-MS analysis of the antennae

The antennae samples (with 6 biological replicates in each group) were homogenized with liquid nitrogen. Then, each sample was sufficiently vortexed for 1 min using pre-cooled 50% methanol of 120 µL, followed by 10-min incubation at room temperature. After overnight storage at −20 °C, the extraction mixture was centrifugated for 20 min under 4,000 *g*, followed by transferring the supernatant into another 96-well plate. Besides, each extraction mixture of 10 μL was combined to prepare pooled quality control (QC) samples [28].

An Ultimate 3000 HPLC (Thermo Scientific) was used to carry out all chromatographic separations. RP separations were realized by adopting the ACQUITY UPLC BEH C18 column (100 mm × 2.1 mm, 1.8 µm, Waters), maintaining the column oven at 50°C. Flowing at 0.3 mL/min, the mobile phase contained phase A (0.1% FA in H_2_O) and phase B (0.1% FA in ACN). Each sample of 4 µL was injected. First scan one or two WASH samples, followed by 3-4 QCs, then insert a QC for every 10 samples scanned, and finally insert 2 QCs. The first and secondary order spectrum data of the metabolites eluted from the column were collected using a Q-Exactive tandem mass spectrometer of high resolution (Thermo Scientific) which was run in both ion modes (negative, NEG, and positive, POS). In the whole acquisition process, a QC sample was taken every after 10 samples to assess LC-MS stability.

### 2.7 Analysis of metabolomics data

MS data were pre-processed by importing the MS data in Compound Discoverer 3.1.0 (Thermo Scientific). This involves extracting peaks, correcting intra-group and inter-group retention time, merging adduct ions, filling gaps, labeling background peaks and identifying metabolites. By applying online databases KEGG and Human Metabolome Database (HMDB), metabolites were annotated through matching to data about exact molecular mass, sample name and formula with those in the databases. The metabolite identification was validated by adopting an in-house fragment spectral library for metabolites.

The metaX was applied to further pre-process the intensity of peak data [29]. After removing features detected in 80% of biological samples or less than 50% of QC samples, the k-nearest neighbor algorithm was used to input the remaining peaks, the values of which were missed, for further improving data quality. Outliers were detected and batch effects were assessed through principal-component analysis (PCA) through use of the pre-processed dataset. The data were normalized by using the probabilistic quotient normalization method, so as to obtain normalized ion intensity data of each sample. Robust LOESS signal correction based on QC was fitted with QC data in terms of the injection order for the purpose of minimizing drift of signal intensity with time. Besides, after calculating relative standard deviations for metabolic features of every QC samples, those larger than 30% were removed.

### 2.8 Statistical analysis of the DEMs

For screening of the DEMs, metaX was used to perform supervised learning method partial least squares-discriminant analysis (PLS-DA), for discriminating disparate variables across groups. After calculating VIP values, a VIP cut-off value of 1.0 was applied for selection of key features. Based on the student’s t‐test algorithm, FC of mean values of normalized ion intensities for various features across control and experimental groups were solved. KEGG enrichment analysis was performed on significantly different metabolites (FC ≥ 2 or ≤ 0.5, p ≤ 0.05, VIP ≥ 1).

### 2.9 Validation of proteome data by quantitative RT-PCR

After being extracted from experimental and control samples using TRIzol^™^ reagent (15596018, Invitrogen), the total RNA was reversely transcribed into cDNA with the PrimeScript^™^ RT reagent Kit (RR047A, TaRaKa) according to the manufacturers’ protocols. The CDSs of the candidate genes used for qRT-PCR referred to the online database (https://bigd.big.ac.cn/gwh/, AN: GWHABGR00000000) [27]. After being devised using the Primer Premier 5.0 software, primers specific for genes are listed in Table S1. By using a 20 μL reaction system, qRT-PCR was conducted in a CFX Connect^™^ Real-Time System (Bio-Rad) following this process: 95 °C for 30 s, then 95 °C for 5 s for 39 cycles, and finally 60 °C for 30 s. After each run, melting curves were produced for confirmation of individual PCR products. All reactions were operated for each time. The amplified transcripts was quantified by *C*t value of each transcript [30] and normalized to the housekeeping gene *Apactin1* (GenBank: GKC242321.1) of *A. pernyi*.

## 3. Results

### 3.1 Identification of the DEPs from the WF male antennae compared with control

Based on the criteria that the quantification of proteins with a ratio (the WF/control male antennae) ≥ 1.2 or ≤ 1/1.2 (p ≤ 0.05), a total of 92 DEPs in the WF group, including 52 up-regulated and 40 down-regulated proteins (Fig. 1A, Table S2), were identified when comparing to the control. Half of the DEPs were defined by more than two peptides (Fig. 1B), and the molecular weights (Mw) of the DEPs mainly ranged from 10 and 80 kDa (Fig. 1C). Statistical analysis of the above proteins showed that they have adequate sequence coverage: 48.9% of these proteins with sequence coverage larger than 10%; 34.8% identified with sequence coverage from 10 to 20% (Fig. 1D). The DEPs were hierarchically clustered for the purpose of determining expression patterns (Fig. 1E).

**Fig. 1.**
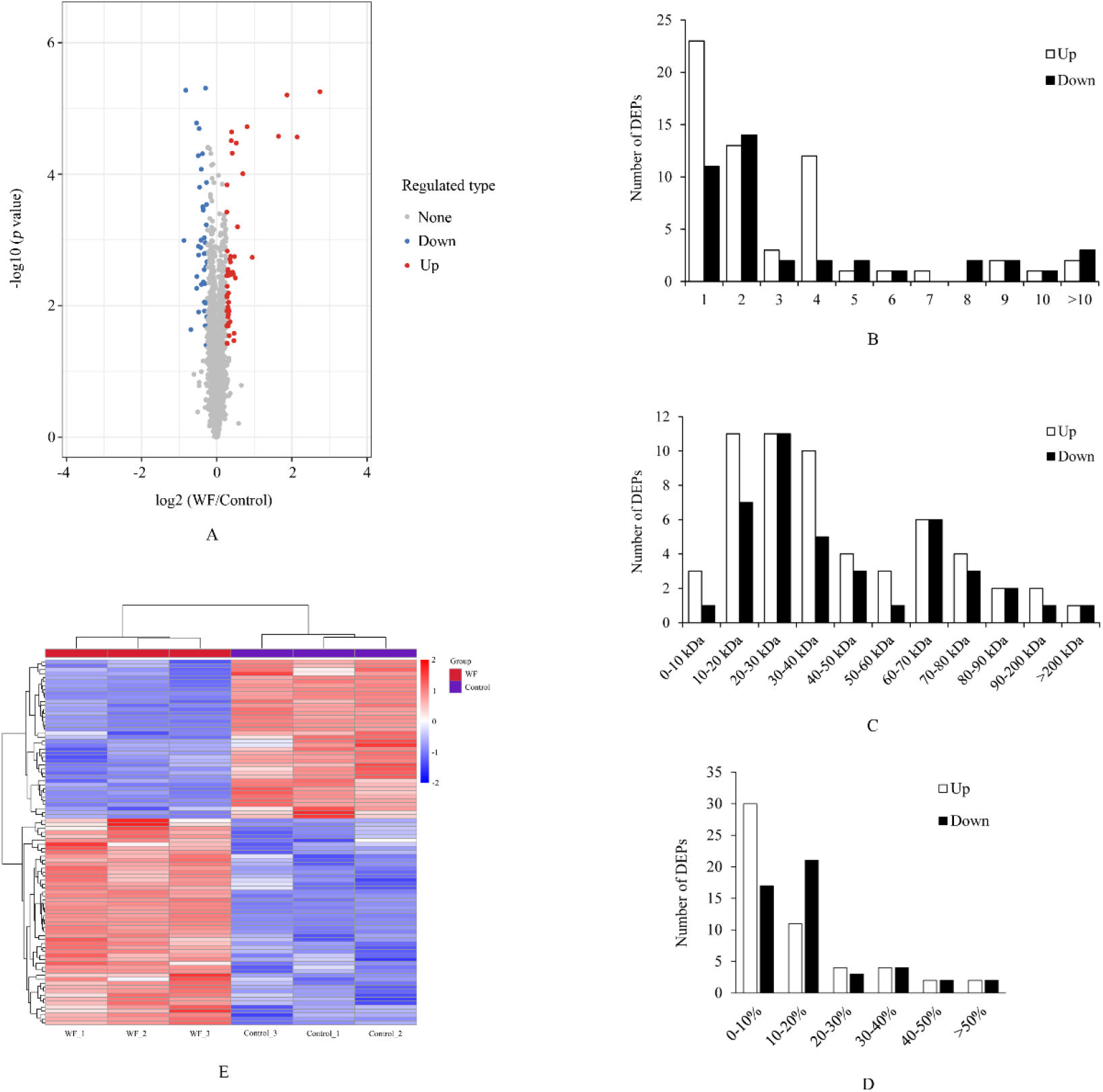
Basic information of the DEPs on the TMT-based proteomics analysis. (A) Volcano plot of the DEPs. The *X-* and *Y*-axis separately denote fold-changes of proteins between the WF and Control groups and the statistical significance of changes in various proteins. Each of dots in this volcano plot denotes one protein. Red, blue, and grey dots separately represent proteins that are significantly up-regulated, down-regulated, and not regulated. (B) Distribution of the peptide number in each DEP. (C) Mass distribution of the DEPs. (D) The sequence coverage distribution of the DEPs. (E) Hierarchical clustering for the DEPs between the control and WF groups. Each column represents each of the samples. The various colors indicate disparate protein expression levels. The protein expression level declines as the color changes from red to blue.

### 3.2 Bioinformatics analysis of the DEPs

GO classification was conducted for annotating the DEPs. Totally 45 DEPs (18 up-regulated and 27 down-regulated) were ascribed to biological process, molecular function, and cellular component at GO terms level 1, and 27 categories at GO terms level 2 (Fig. 2A, Table S3). Dominant GO terms for subcategories of biological process were classified into the biological regulation, as well as cellular, multicellular organismal, developmental, and metabolic process. Subcategories of cellular component were mainly assigned to cell, intracellular, and protein-containing complex. The largest subcategories of molecular function were binding and catalytic activity.

**Fig. 2.**
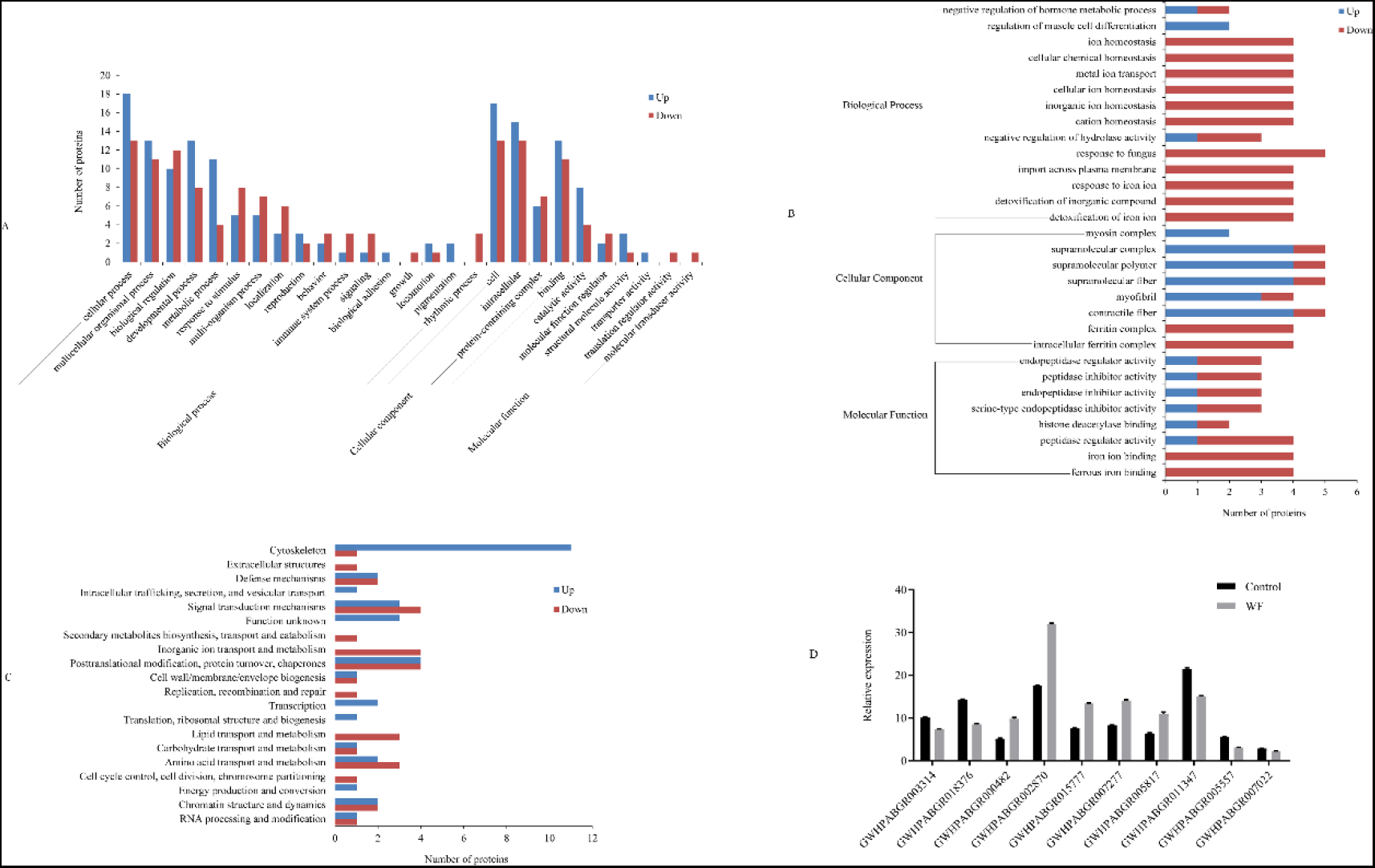
Bioinformatics analysis including GO classification (A), GO enrichment analysis (B) and COG classification (C) of the DEPs, and validation of the DEPs selected via qPCR. (D). Red and blue bars indicate up-regulated and down-regulated proteins respectively. The *X*- and *Y*-axis in (D) separately represent the selected proteins for qPCR and relative expressions. GWHPABGR003314, neuroendocrine convertase 1-like. GWHPABGR018376, ferritin. GWHPABGR000482, tyrosine hydroxylase. GWHPABGR002870, cryptochrome-1. GWHPABGR015777, tachykinin. GWHPABGR007277, arylalkylamine *N*-acetyltransferase. GWHPABGR005817, cadherin-23. GWHPABGR011347, glutathione *S*-transferase delta 3. GWHPABGR005557, peptidoglycan-recognition protein. GWHPABGR007022, acyl-peptide hydrolase.

Further GO enrichment analyses revealed enrichment of the DEPs into 30 GO terms (Fig. 2B, Table S4). Under the biological process category, detoxification of iron ion, detoxification of inorganic compound, response to iron ion, import across plasma membrane, response to fungus, cation homeostasis, inorganic ion homeostasis, cellular ion homeostasis, metal ion transport, cellular chemical homeostasis, and ion homeostasis were significantly enriched, and the mapped DEPs were mainly down-regulated. Besides, under cellular component and molecular function, terms for intracellular ferritin complex, ferritin complex, contractile fiber, myofibril, and ferrous iron binding, iron ion binding, peptidase regulator activity were remarkably enriched. The changes of expression levels of the proteins related to ion metabolism may suggest the changes in transmission of neural signal in male antennae following the stimulation of females.

COG is a database in which orthologous proteins are classified, providing information about the possible functions of the DEPs. COG analysis showed that 61 DEPs were assigned to 20 functional categories (Fig. 2C, Table S5). The main categories were cytoskeleton, posttranslational modification, protein turnover, chaperones, and signal transduction mechanisms, followed by amino acid transport and metabolism, defense mechanisms, inorganic ion transport and metabolism, chromatin structure and dynamics, and lipid transport and metabolism.

### 3.3 qRT-PCR verification of the DEPs

In a bid to verify that proteome data are accurate, we selected 10 DEPs including 5 up- and 5 down-regulated proteins (Table S1), to carry out qRT-PCR analyses for verifying their transcriptional expression. The results also show agreement of candidate genes with those in the proteome in terms of expression patterns (Fig. 2D), which verifies DEP expression in deep sequencing analyses.

### 3.4 Metabolome data summary

The total ion chromatograms of features determined in antennae from the control and WF groups in NEG and POS modes showed that the six repeated biological samples of the control and experimental groups are found to be highly coincident, respectively, which indicates biological replicates to be insignificantly different (Fig. S1). Totally 6343 and 5957 features containing 6157 and 5493 high-quality (HQ) ones were separately produced in the NEG and POS modes (Table 1). Meanwhile, 1210 DE features including 716 up-regulated and 494 down-regulated features were screened via the multiple statistical analysis based on metabolomic profiling (Table 2, Fig. 3A). This was followed by annotation of the DEMs through matching the DE features with KEGG and HMDB databases and verification of them on the basis of comparing spectra for secondary fragmental ions by means of MS/MS. In this way, 545 DEMs covering 218 up-regulated and 327 down-regulated ones were identified from the WF group compared with control (Fig. 3B & C, Table S6).

**Table 1.** Statistical analysis of quantitative features from the metabolomic data

| Index | POS | NEG | Total |
| --- | --- | --- | --- |
| All features | 5957 | 6343 | 12300 |
| HQ features | 5493 | 6157 | 11650 |
| DE features | 603 | 607 | 1210 |

**Table 2.**
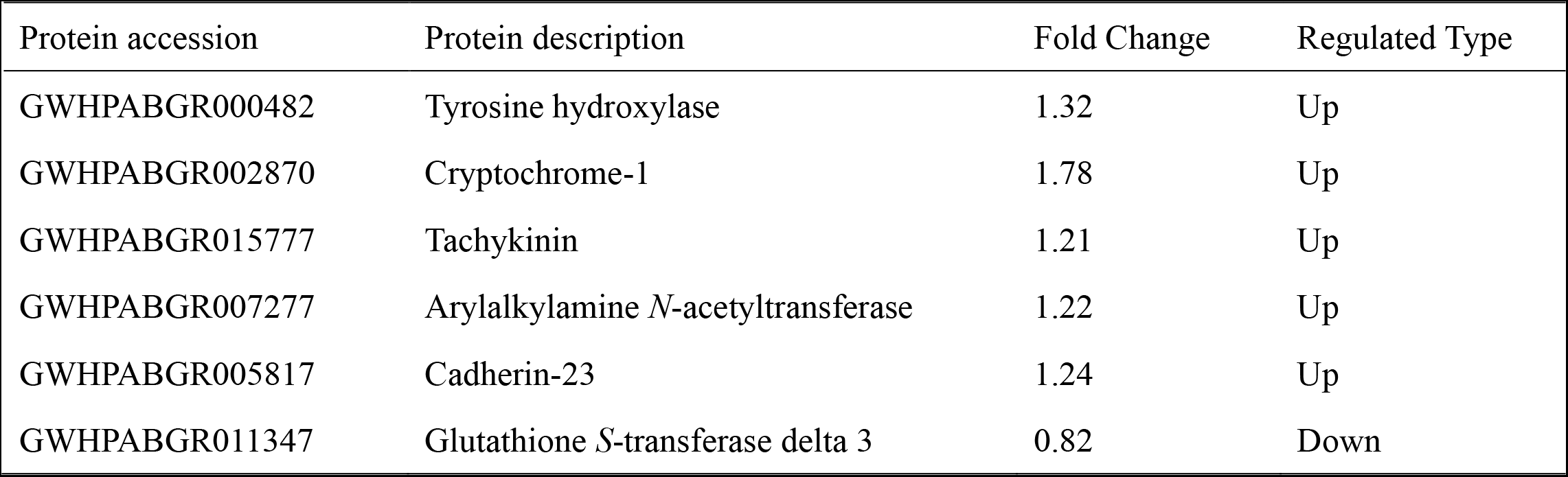
The candidate DEPs involved in neural transmission of female pheromone signals

| Protein accession | Protein description | Fold Change | Regulated Type |
| --- | --- | --- | --- |
| GWHPABGR000482 | Tyrosine hydroxylase | 1.32 | Up |
| GWHPABGR002870 | Cryptochrome-1 | 1.78 | Up |
| GWHPABGR015777 | Tachykinin | 1.21 | Up |
| GWHPABGR007277 | Arylalkylamine *N*-acetyltransferase | 1.22 | Up |
| GWHPABGR005817 | Cadherin-23 | 1.24 | Up |
| GWHPABGR011347 | Glutathione *S*-transferase delta 3 | 0.82 | Down |

**Fig. 3.**
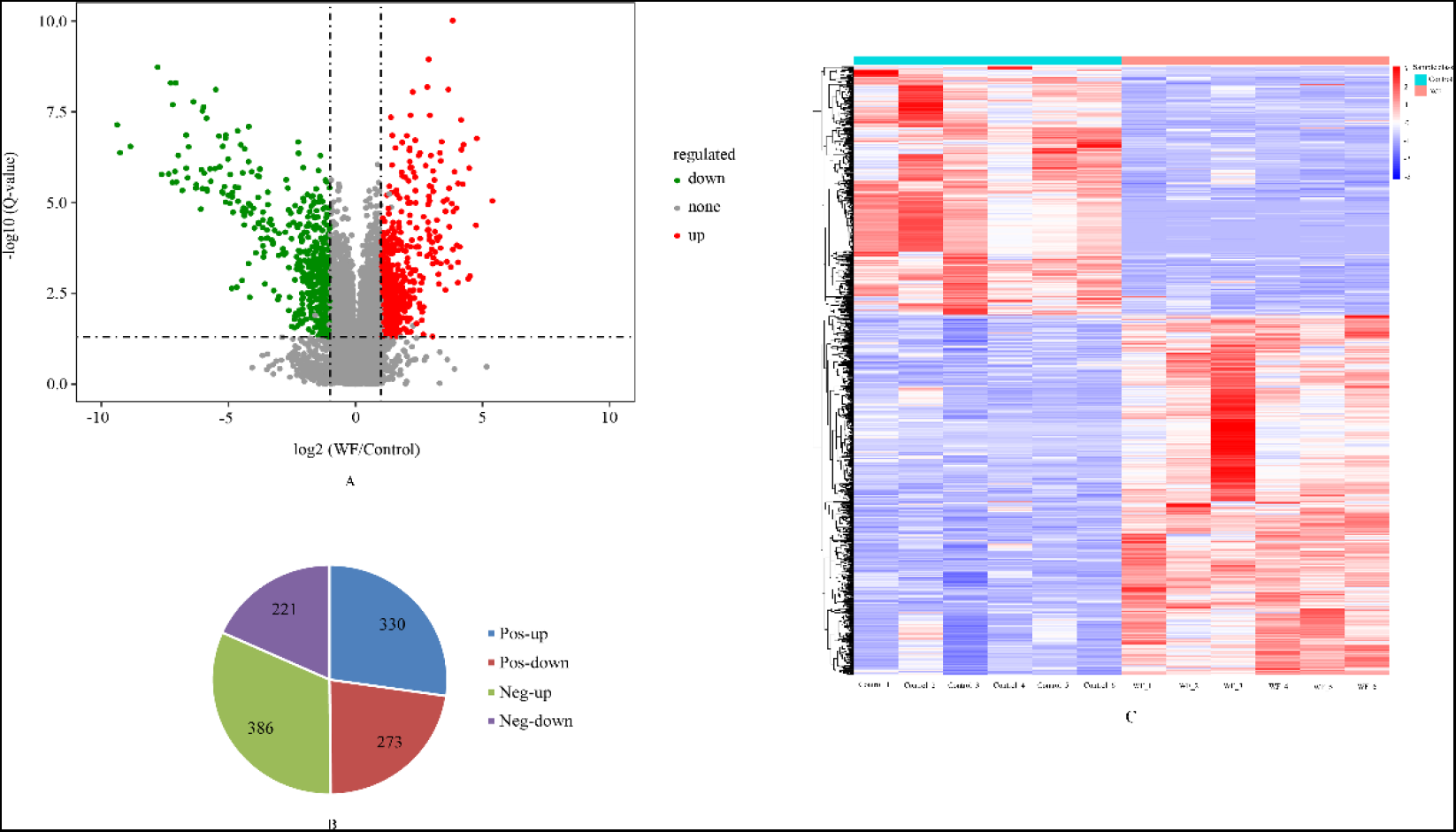
Statistical charts for the DE features and DEMs. (A) Volcano plot for the DE features of the WF group in comparison with the control. The *X-* and *Y*-axis separately represent fold-changes of features across various experimental groups and statistical significance of changes in various features. Each of dots denotes one feature. Red and blue dots separately indicate features that are significantly up-regulated and down-regulated. (B) Numbers of DE features in the both ion modes (POS and NEG). (C) Hierarchical clustering for the DEMs between the control and WF groups. Each column represents each of the samples. The different colors indicate disparate metabolite expression levels. The metabolite expression level declines as the color changes from red to blue.

The PCA analysis of the DEMs indicated good inter-separation and intra-clustering of samples of the WF and control groups (Fig. S2A & B), implying that each group to be highly metabolically repeatable and the two groups to show high metabolic diversity. For further confirming metabolic variation of samples in the WF and control groups, the PLS-DA models were run, revealing that the two groups exhibited significant metabolic differences (Fig. S2C & D). The 200 permutation tests verified the PLS-DA models to be credible and non-overfitted (Fig. S2E & F).

### 3.5 Classification and KEGG pathway enrichment analysis of the DEMs

According to HMDB annotation, a total of 160 DEMs (88 up-regulated and 72 down-regulated) were classified into 11 and 44 categories on SuperClass and Class level, respectively (Fig. 4, Table S6). Among the Class level classifications, the DEMs were mainly grouped into benzene and substituted derivatives, fatty acyls, carboxylic acids and derivatives, and organooxygen compounds. Attaching fatty acyl groups as thioesters to the internal cysteine contributes the unique reversible protein lipidation form, protein *S*-acylation, also known as palmitoylation of proteins playing a crucial part in diverse physiological processes like Ras signaling, intracellular trafficking, neuronal scaffolding protein localization, and ion channel activity [31–37]. The metabolic changes of certain DE fatty acyls such as down-regulation of palmitamide may decline the S-acylation level thus affect the signal transduction in neurons. Among the carboxylic acid metabolites, the up-regulation of fumaric acid which is one of the intermediate metabolites of the tricarboxylic acid cycle may reflect the increase of energy metabolism for neurotransmission.

**Fig. 4.**
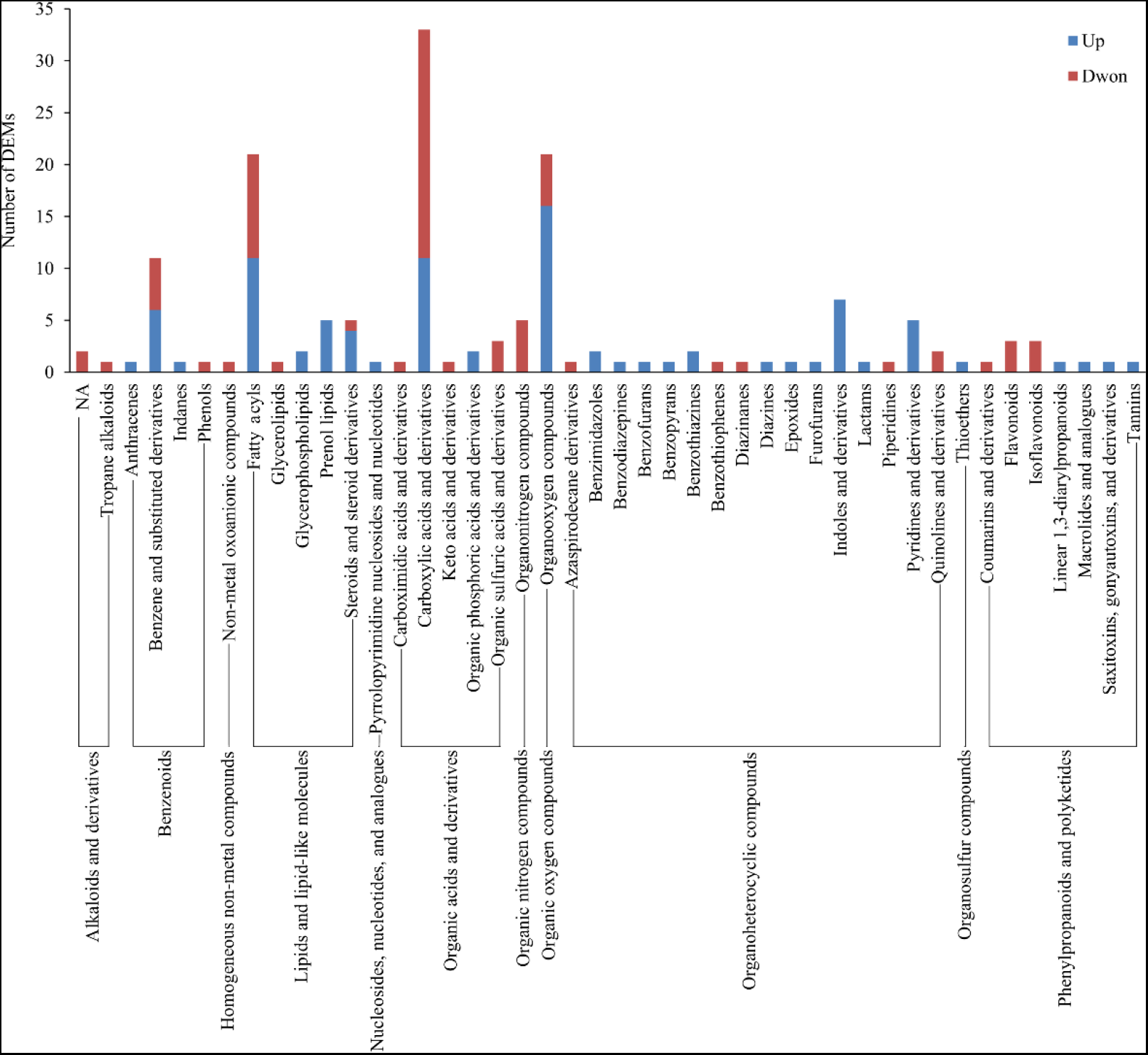
Classification of the DEMs based on HMDB annotation. The *X-* and *Y*-axis separately indicate the DEM number and the categories of the DEMs at SuperClass and Class levels.

KEGG enrichment analysis indicated enrichment of totally 43 DEMs (26 up-regulated and 17 down-regulated) into 6, 27, and 87 pathways at level 1, 2, and 3, respectively (Fig. 5, Table S6). The DEMs were mainly enriched into metabolism-related pathways, such as metabolism of amino acid, carbohydrate, energy, lipid, nucleotide, cofactors, and vitamins, xenobiotics biodegradation and metabolism, biosynthesis of other secondary metabolites. Several pathways involved in environmental information processing were enriched, including phosphotransferase system, ABC transporters, sphingolipid signaling pathway, neuroactive ligand-receptor interaction, implying their responsibilities underlying female pheromone processing in male antennae following female stimulation.

**Fig. 5.**
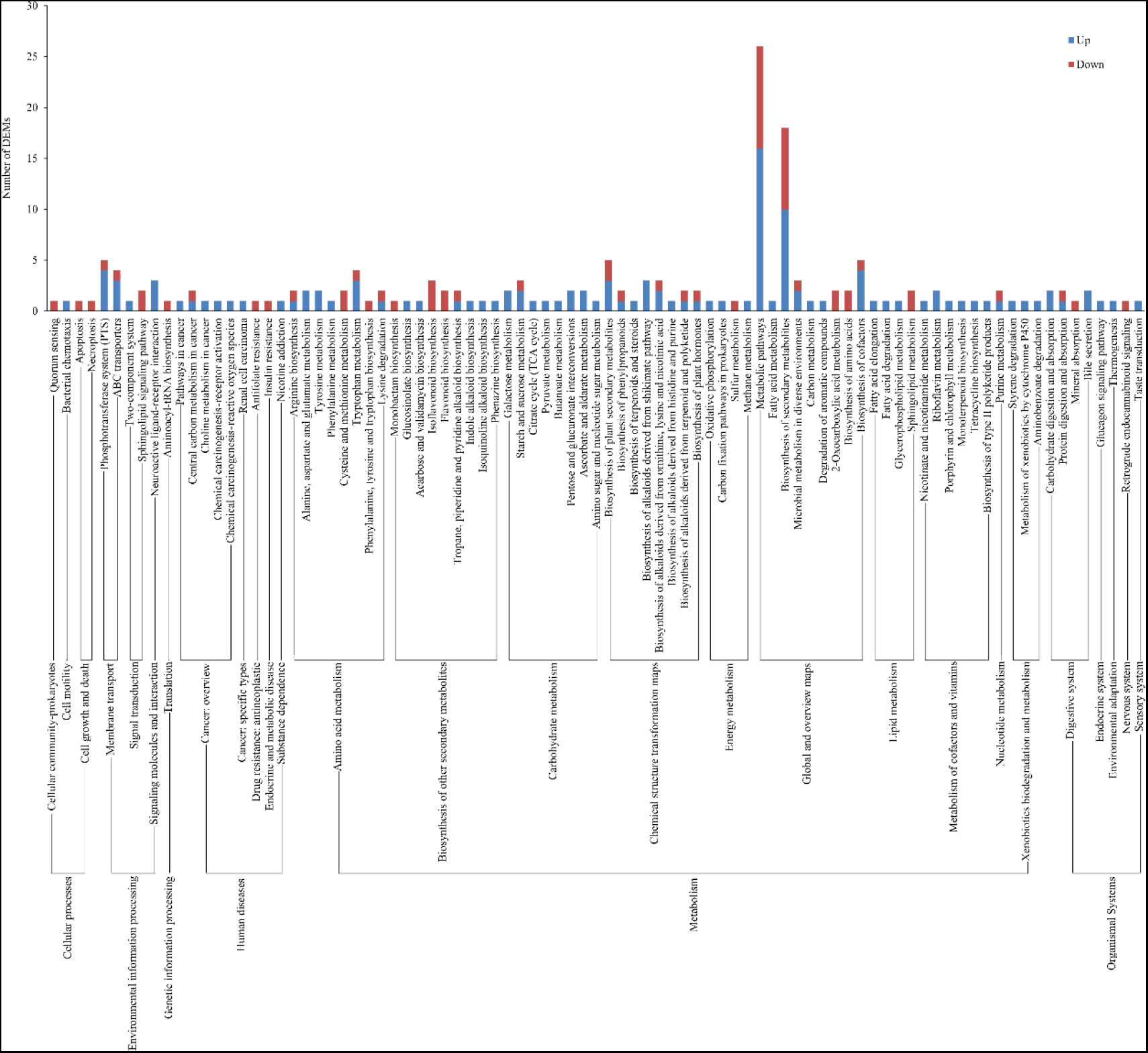
KEGG pathway enrichment analysis for the DEMs. The metabolic pathway categories at level 1, 2, and 3 are indicated by the *X*-axis while the DEM number is shown on the *Y*-axis.

## 4. Discussion

Before mating process, the *A. pernyi* moths require a series of information exchanges between males and females. After receive the female pheromones, the male antennae exhibit certain physiological responses which gradually stretching out from clinging to the head (Fig. 7). Then the male moths initiate courtship behavior which represent as wing-flapping. Reception of the sex pheromone via antennae is fundamental for the males to court and mate with females. In this study, to enable male moths to perceive the female sex pheromone adequately, the males and females were contact with each other for 2-5 min until each male moth exhibited wing-flapping. Integrative proteomics and metabolomics analysis were carried out to compare the proteins and metabolites in the antennae that were differentially expressed between the wing-flapping and control males for exploring the response mechanisms of the male antennae following stimulation by the female sex pheromone, and we mainly characterized the DEPs and DEMs involved in neural transmission of sex pheromone signals in the male antennae.

Based on the proteomic analysis, 6 DEPs involved in regulating signal transduction were screened, including tyrosine hydroxylase (TH, GWHPABGR000482), tachykinin (TK, GWHPABGR015777), cryptochrome-1 (CRY-1, GWHPABGR002870), arylalkylamine *N*-acetyltransferase (AANAT, GWHPABGR007277), cadherin-23 (Cdh-23, GWHPABGR005817), and glutathione *S*-transferase delta 3 (GST-δ3, GWHPABGR011347) (Fig. 6A, Table 2). Insect GSTs of are classified into several subgroups (microsomal, cytosolic, and mitochondrial) according to the cellular location [38]. Most cytosolic GSTs of insects fall in six subclasses including δ, ε, ω, θ, σ, and ζ [39, 40] with δ and ε are insect-specific [41]. Meanwhile, a series of antennae-specific GSTs which function as scavengers of pheromones and host volatiles in the odorant detection system have been identified in several insect species such as *Manduca sexta*, *Papilio xuthus*, and *Plodia interpunctella* [42–44]. In *Bombyx mori*, BmGSTd4 which is a GST specific for antennae in male moths has been demonstrated possessing dual functions in detoxifying xenobiotic compound and terminating signals of sex pheromone [45]. We found that in the antennae of male *A. pernyi*, GST-δ3 was down-regulated after being stimulated by females, which may suggesting its negative regulation in continuous reception of female pheromones by the male antennae.

**Fig. 6.**
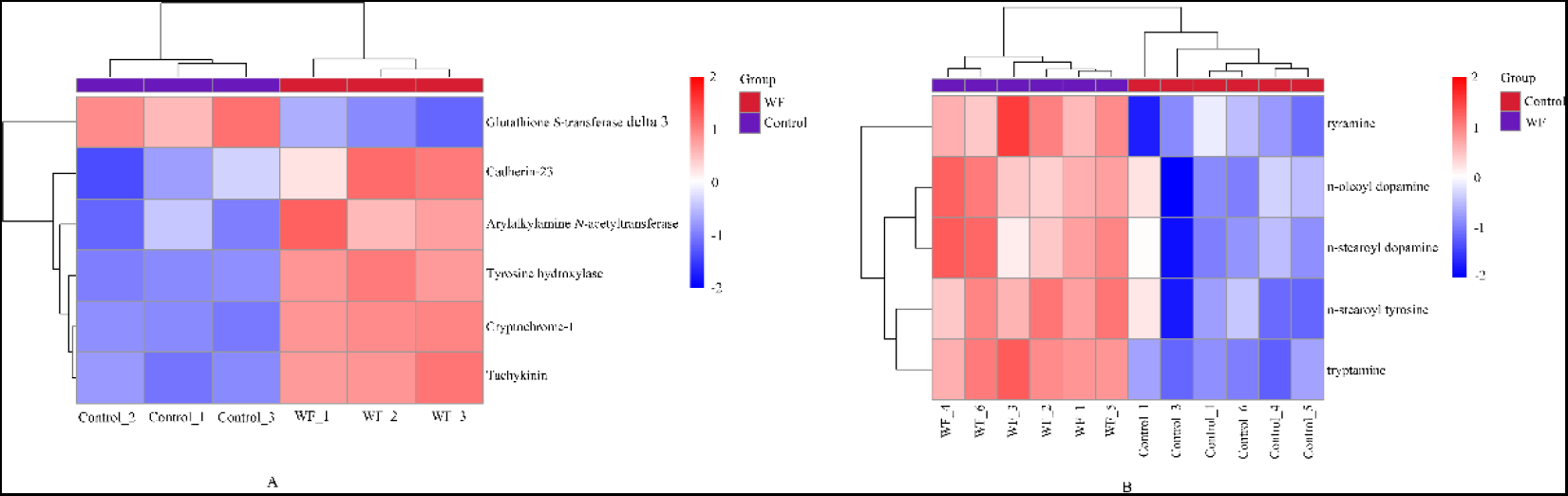
Hierarchical clustering of the candidate DEPs (A) and DEMs (B) involved in neural transmission of female pheromone signals between the control and WF groups. Different columns represent different samples. Different colors represent different levels of protein expression. The protein expression levels decline as color changes from red to blue.

TH is the rate-limiting enzyme for synthesizing catecholamines, including dopamine (DOPA), epinephrine and norepinephrine, which serve as important neurotransmitters and hormones in peripheral and central nervous systems and are products of this cascade pathway [46]. Once catecholaminergic synapses need more neurotransmitters, TH is activated for producing more DOPA which is transferred into synaptic vesicles following decarboxylation with continuous synthesis of other catecholamines. DOPA and several types of dopaminergic neurons have been demonstrated that are involved in regulating olfactory memory formation, strength and stability, and promoting motor activity and arousal in a variety of insects such as *Drosophila melanogaster* and *Apis mellifera* [47–55]. The depletion of DOPA induced by TH inactivation decreased arousal state and motor activity in *D*. *melanogaster* [56]. In addition, DOPA could be acetylated by AANAT, for which *N*-acetylation is regarded as the main way of transformation of biogenic amines, in the nervous system of insects [57]. In this study, the up-regulation of TH and AANAT may suggest an increasement of DOPA for increasing transmission of the female pheromone signals from the antennae to the brain of male *A. pernyi*, thus leading to the motor activity enhancing of the male moths.

Cdhs in the central nervous system of vertebrates serve as primary organizers to form and remodel the synapse, and associate with intracellular binding proteins to form cadherin adhesion complexes for modifying adhesive contact, regulating synapse formation and plasticity, and activating signal transducing pathway [58, 59]. As a member of the Cdh family, the up-regulation of Cdh-23 may be involved in activation of the female pheromone signals transmission between cells in the antennae of male *A. pernyi*. CRYs serve as a circadian clock component for controlling daily behavioral and physiological rhythms and a photoreceptor for mediating light entrainment of insect circadian clock [60]. Recently, CRY has been proved to regulate the courtship activity increasing in a magnetic field of male *D*. *melanogaster* which effect was disrupted due to CRY-deficient and RNAi-mediated knockdown of *cry* [61]. TK is a kind of multifunctional conserved neuropeptide and play important roles in nerve transmission or as neuromodulator in the central and peripheral nervous system. TKs also have been proved to be involved in regulating locomotor activity and olfactory perception [62, 63]. In *D*. *melanogaster*, silencing TK-related peptides caused perception for specific concentrations and odorants to change in a trend to lose sensitivity, therefore leading to substantial changes in behavioral responses to indifference [62]. In this study, it is possible that the up-regulations of CRY-1 and TK were related to the increased motor activity of the male antennae for searching and precise positioning the female pheromones in the environment and the hence wing-flapping activity.

From the metabolome, we screened 2 biogenic amines including tyramine (TyrA) and tryptamine (TryA) which were differentially expressed in the antennae of male *A. pernyi*. The nervous system of animal produces coordinated, integrated behaviors via enabling communication related to speed of delivery, privacy, and duration of the message to be flexible by adopting plenty of neuroactive chemicals that act as neurotransmitters, neurohormones, or neuromodulators [64]. It has been demonstrated that tyramine obtained via synthesis by tyrosine decarboxylase using tyrosine capable of being released from neurons belongs to the predominantly neuroactive chemicals of insects [65]. Tryptamine is a serotonin-related indolamine which is involved in regulating interactions between serotoninergic and catecholaminergic systems [66]. Besides, 3 DEMs involved in neuroprotection, including *N*-stearoyl tyrosine (NsTyr) which is an anandamide analogue and could protect neurons from apoptosis [67], *N*-oleoyl dopamine (NODA) and *N*-stearoyl dopamine (NSDA) belonging to *N*-acyl dopamine possessing neuroprotective activities [68, 69], were screened. We concluded that the up-regulation of those DEMs (Fig. 6B, Table 3) may also contribute to the enhanced transmission of the neural impulses between neurons in the male antennae via regulating the neuron activity.

**Table 3.** The candidate DEMs involved in neural transmission of female pheromone signals

| Metabolic ion ID | Metabolite description | Fold Change | Regulated Type |
| --- | --- | --- | --- |
| pos-3.681_137.08401 | Tyramine | 2.37 | Up |
| pos-3.42_160.10008 | Tryptamine | 7.61 | Up |
| neg-7.642_417.3247 | *N*-oleoyl dopamine | 2.14 | Up |
| neg-8.323_419.34033 | *N*-stearoyl dopamine | 2.43 | Up |
| neg-7.876_447.33527 | *N*-stearoyl tyrosine | 2.23 | Up |

The transmission of female pheromone signals in male antennae is a complex biological process. Based on integrated proteomics and metabolomics data of the current study, a presumptive model elucidating response mechanisms for the male moth antennae of *A. pernyi* to female pheromones was displayed in Fig. 7. On one hand, several factors were involved in regulating the neuron activity for enhanced transmission of neural impulses, including the up-regulation of NODA and NSDA generated via *N*-acylation of DOPA which production and *N*-acetylation was elevated due to the up-regulated TH and AANAT receptively, and the increased expression level of TyrA, TryA, and NsTyr produced by *N*-acetylation of tyrosine. On the other hand, the down-regulation of GST-δ3 and up-regulation of CRY-1, TK, and Cdh-23 were related to continuous perception, reception, and transduction of female pheromone signals. In general, our findings provide new clues on further exploration of potential molecular mechanisms underlying the courtship behavior of male *A. pernyi*.

**Fig. 7.**
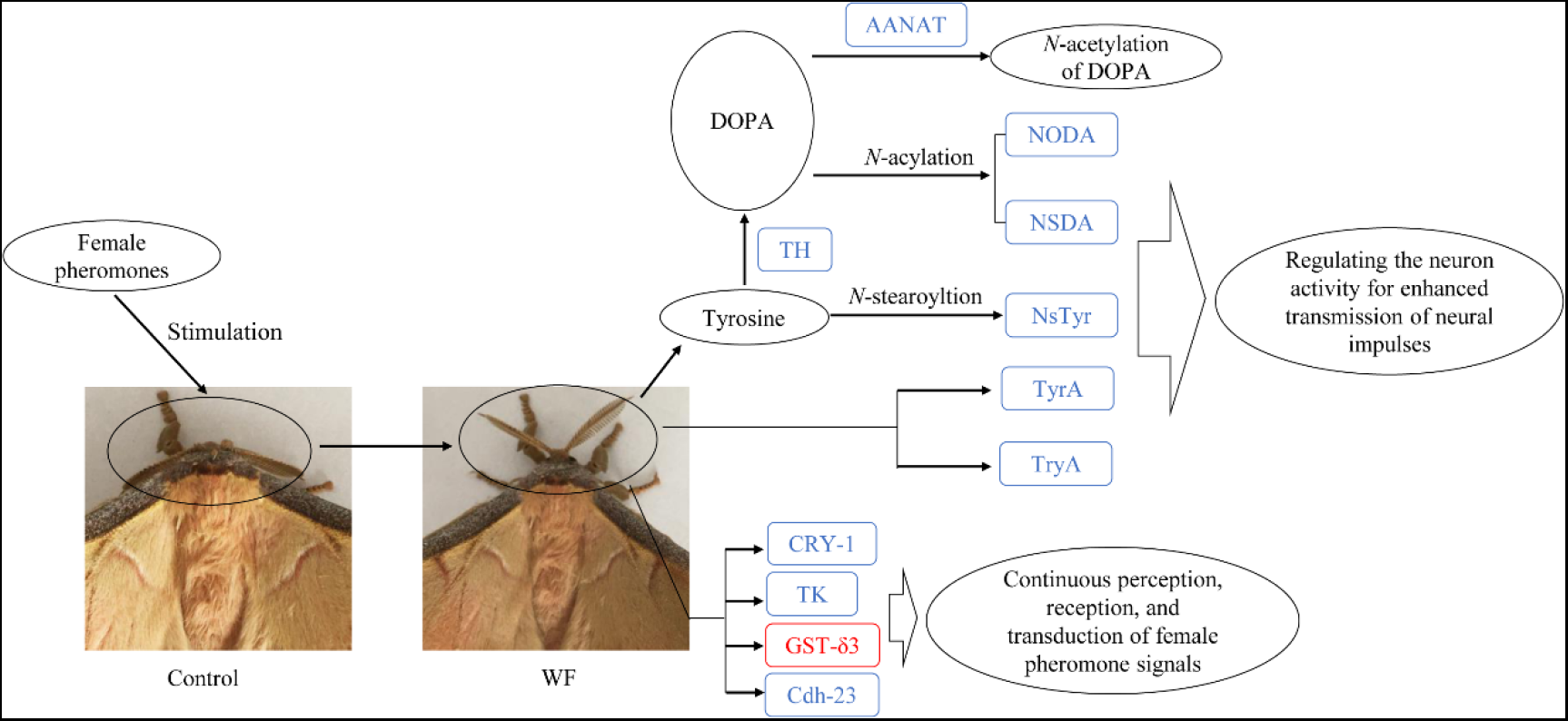
Hypothetical model of changes in the male moth antennae of *A. pernyi* following female pheromone stimulation. Blue (red) fonts denote up-regulated (down-regulated) proteins or metabolites.

## Acknowledgements

This work was supported by the Natural Science Foundation of Shandong Province (ZR2020QC190), Weifang University Doctor Startup Fund (2020BS26), and Agricultural Science and Technology Innovation Project of Shandong Academy of Agricultural Sciences (CXGC2022B02).

